# Viscous medium-based crystal support in sample holder for fixed-target serial femtosecond crystallography

**DOI:** 10.1101/2019.12.31.892265

**Authors:** Keondo Lee, Donghyeon Lee, Sangwon Baek, Jaehyun Park, Sang Jae Lee, Sehan Park, Wan Kyun Chung, Jong-Lam Lee, Hyun-Soo Cho, Yunje Cho, Ki Hyun Nam

**Author notes:** Keondo Lee, Donghyeon Lee and Sangwon Baek contributed equally. Synopsis We developed a viscous medium-based crystal support in a sample holder for fixed-target serial crystallography.

## Abstract

Serial femtosecond crystallography (SFX) enables the determination of a room-temperature crystal structure of macromolecules without causing radiation damage, as well as provides time-resolved molecular dynamics data in pump-probe experiments. Fixed-target SFX (FT-SFX) can minimize sample consumption and physical effects to crystals during sample delivery. Various types of sample holders have been developed and applied in FT-SFX; however, no sample holder has been developed that can universally mount crystals of various sizes and shapes. Here, we introduce a viscous media-based crystal support in a sample holder for FT-SFX. Crystal samples were embedded in viscous media such as gelatin and agarose, which were enclosed in a polyimide film. In the vertically placed sample holder, the viscous medium stably supported crystals between the two polyimide films without crystal sinking due to gravity. Using this method, we performed FT-SFX experiments with glucose isomerase and lysozyme embedded in gelatin and agarose, respectively. The room-temperature crystal structures of glucose isomerase and lysozyme were successfully determined at 1.75 and 1.80 Å resolutions, respectively. Viscous media used in this experiment showed negligible background scattering in data processing. This method is useful for delivering crystal samples of various sizes and shapes in FT-SFX experiments.

## 1. Introduction

Serial femtosecond crystallography (SFX) can visualize the time-resolved molecular dynamics of macromolecules or small molecules at room temperature without causing radiation damage through pump-probe experiments (Chapman *et al.*, 2014, Nogly *et al.*, 2015, Weinert *et al.*, 2017, Martin-Garcia *et al.*, 2017). SFX using an intense X-ray free electron laser (XFEL) with an ultrashort pulse can be used to analyze smaller crystals than those used in traditional X-ray crystallographic methods; however, this method requires a large number of crystal samples, as each crystal is exposed only once to the X-ray beam (Boutet *et al.*, 2012). Thus, this technique requires a delivery technology that serially and reliably delivers numerous crystal samples to the X-ray interaction position (Grunbein & Nass Kovacs, 2019). Various sample delivery methods such as injector-based methods (DePonte *et al.*, 2008, Weierstall *et al.*, 2014), syringe method (Sugahara *et al.*, 2015, Berntsen *et al.*, 2019, Park & Nam, 2019), capillary method (Stellato *et al.*, 2014, Nam, 2020a), fixed-target scanning (Hunter *et al.*, 2014, Cohen *et al.*, 2014, Mueller *et al.*, 2015, Lee *et al.*, 2019), electrospinning (Sierra *et al.*, 2012), and microfluidic devices (Monteiro *et al.*, 2019) have been used for crystal delivery in serial crystallography (SX) using XFEL or synchrotron X-rays.

Among them, fixed-target (FT) scanning is widely used in both traditional and serial crystallography (SX) studies (Rodgers, 1994, Hunter *et al.*, 2014, Cohen *et al.*, 2014, Mueller *et al.*, 2015, Roedig *et al.*, 2017, Lee *et al.*, 2019). In FT-SFX, a large number of crystal samples are supported in a sample holder; data collection involves several components, such as the motion stage, goniometer, and conveyor belt (Hunter *et al.*, 2014, Cohen *et al.*, 2014, Mueller *et al.*, 2015, Fuller *et al.*, 2017, Lee *et al.*, 2019, Martiel *et al.*, 2019). This method has the advantages of lower sample consumption and a high hit rate compared to other delivery methods (Oghbaey *et al.*, 2016). Moreover, the injector delivery method may physically affect the crystal sample when it is physically pushed to the X-ray beam path (Martiel *et al.*, 2019). In contrast, in the FT scanning method, since the placed sample holder is continuously provided to the X-ray beam path using a driving device, the sample is not physically damaged (Martiel *et al.*, 2019).

To mount a large number of crystal samples on a single chip, sample holders have been fabricated as silicon chips (Murray *et al.*, 2015), silicon nitride (Hunter *et al.*, 2014), microfluidic chips (Lyubimov *et al.*, 2015), and microgrids (Cohen *et al.*, 2014). These sample holders have been successfully applied in FT-SFX studies, but effort is required to ensure precise alignment of the XFEL and sample holder. We previously developed a nylon-mesh based sample holder enclosed with a polyimide film (Lee *et al.*, 2019). Because this holder was composed of X-ray-transparent material, the approach can be applied without precious alignment between the XFEL beam and sample holder.

On the other hand, these crystal sample holders are regularly arranged with holes or pores of the same shape and size. However, crystal size and shape can vary, and it is difficult to redesign and manufacture a sample holder with holes optimized for a specific crystal sample. Although a nylon mesh-based sample holder is useful for selecting the desired size of nylon to match the crystal shape or size, sample loading is difficult when these factors are random. In contrast, a method for dispersing crystal samples between two sheets of mylar film without a sample holder has also been used for FT-SFX (Doak *et al.*, 2018). This method can load crystals of various sizes and shapes, but the crystals between films can sink because of gravity, making the crystal sample distribution non-uniform. Although a crystal sample can be fixed when the two films are in close contact, this causes physical damage to the crystal sample. Therefore, sample holders that are universally applicable to samples of various shapes and sizes are needed.

Viscous medium is widely used for sample delivery in SX experiments and is often used to produce stable injection streams at low speed through an injector or syringe (Weierstall *et al.*, 2014, Sugahara *et al.*, 2015, Conrad *et al.*, 2015, Park *et al.*, 2019, Nam, 2019, Nam, 2020b). In addition, when delivering crystal samples at a low flow rate in a capillary aligned with the X-ray, the viscous medium prevents crystalline samples from sinking (Nam, 2020a). We predicted that viscous medium can be applied to support a large number of crystal samples in sample holder for FT-SFX.

Here we prepared and characterized a viscous material-based crystal support as a sample holding method for FT-SFX. Glucose isomerase and lysozyme were embedded in gelatin and agarose, respectively, and enclosed in polyimide films. The viscous medium prevented the crystalline samples from sinking when the sample holder was placed vertically. The polyimide film prevented dehydration of the crystalline samples enclosed in the viscous medium. Using this approach, we successfully determined the room temperature structures of glucose isomerase and lysozyme at 1.75 and 1.80 Å resolutions, respectively. This approach can be used to mount crystal samples of various sizes and shapes for FT-SFX.

## 2. Methods and materials

### 2.1. Crystallization

Glucose isomerase from *Streptomyces rubiginosus* was purchased from Hampton Research (Cat No. HR7-102, Aliso Viejo, CA, USA). This contains the crystalline in the glucose isomerase solution, which were used directly for FT-SFX experiments without post-crystallization. The crystal size and density of glucose isomerase were 10–100 μm and approximately 3 × 10^8^ crystals/ml, respectively. Lysozyme from chicken egg white was purchased from Sigma-Aldrich (Cat No. L6876, St. Louis, MO, USA). The lysozyme solution (50 mg/ml) was mixed with an equal volume of crystallization solution containing 0.1 M sodium acetate, pH 4.0, 6% (w/v) PEG 8000, and 2.5 M NaCl in a 1.5 ml microcentrifuge tube, followed by incubation at 20°C overnight. The crystal size and density were 15 μm and 5 × 10^7^ crystals/ml, respectively.

### 2.2. Preparation of viscous medium based sample holder

The sample holder consisting of polyimide film was fabricated as previously reported (Lee *et al.*, 2019). Briefly, a 300 μm PVC frame was attached to the 25 μm polyimide film with double-side adhesive polyimide tape (Supplementary Figure 1). Gelatin from bovine skin (G9382) and agarose (AR1095) were purchased from Sigma-Aldrich and GeorgiaChem (Norcross, GA, USA), respectively. Gelatin (15%, w/v) dissolved in crystallization solution in a 1.5 ml microcentrifuge tube was immersed in boiling water (>100°C) for 5 min. Next, 50 μl of melted gelatin solution was transferred to a 100 μl Hamilton syringe (81065) and left to stand until it was solidified. An empty 100 μl Hamilton syringe was connected to a syringe containing the gelatin gel using a coupler. The solidified gelatin was fragmented by mechanical mixing in the dual syringe setup (Nam, 2019, Park *et al.*, 2019). The syringe containing gelatin fragments was connected to a syringe containing 20 μl of the glucose isomerase crystals suspension using a coupler, followed by gentle mixing by moving of the plunger back and forth more than 20 times. The glucose isomerase crystals enclosed in the gelatin fragments were loaded through a needle onto a polyimide film inside the PVC frame (Figure 1). Using a pipette tip, the gelatin fragments containing crystals were gently spread over the entire area of the polyimide film. The samples were covered with a polyimide film using double-side adhesive polyimide tape. Lysozyme crystals were embedded in final 4% (w/v) agarose and loaded onto the polyimide film as described for embedding glucose isomerase in gelatin.

**Figure 1.**
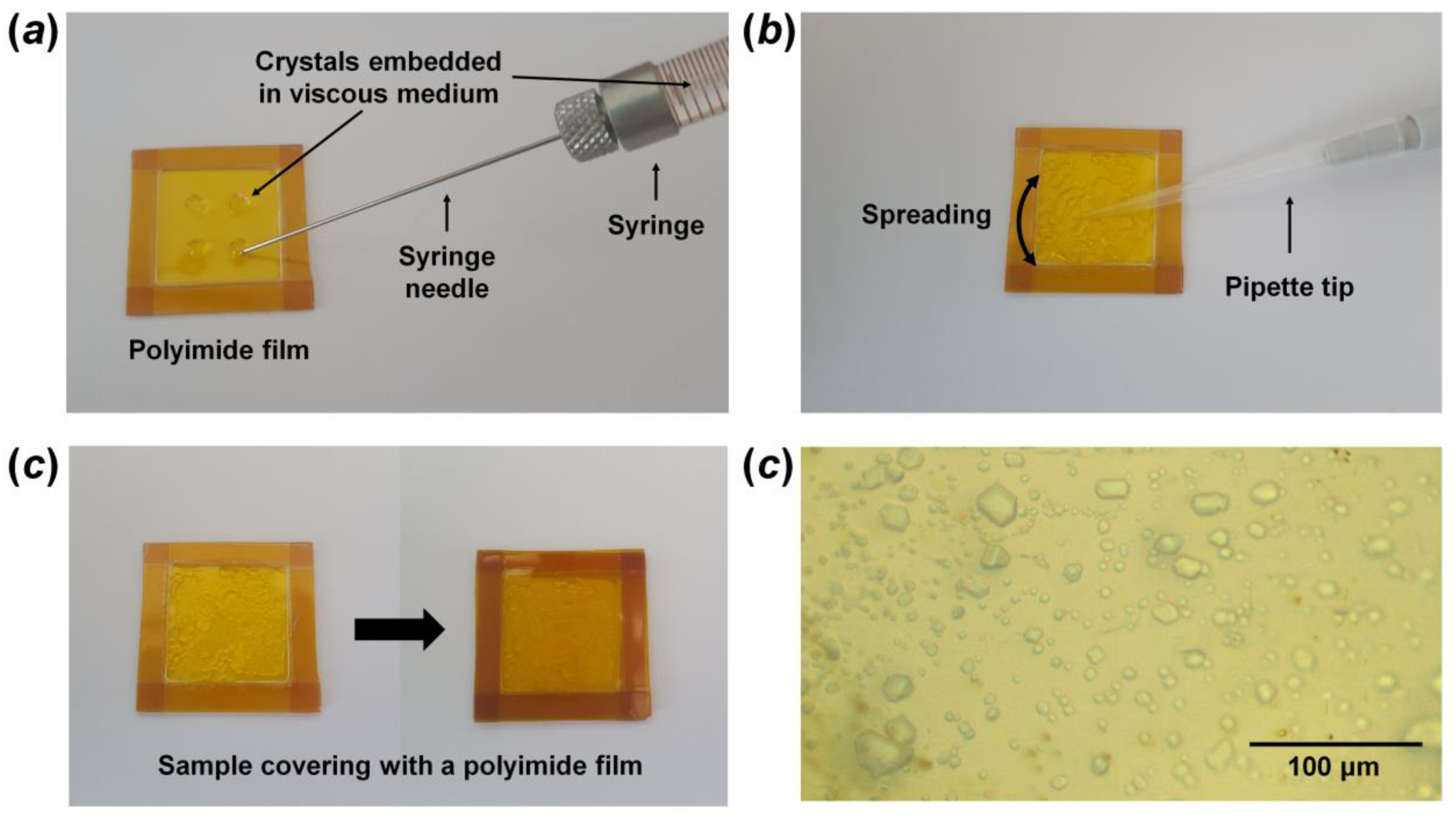
Preparation of viscous medium-based sample holder for FT-SFX. (a) The crystals embedded in gelatin were loaded onto the polyimide chip. (b) The sample was spread evenly using a pipette tip.(c) The polyimide chip was covered to prevent solution evaporation. (d) Microscope view of glucose isomerase embedded in gelatin on the sample holder.

### 2.3. Data collection

FT-SMX was performed at the nano-crystallography and coherent imaging experimental hutch in the Pohang Accelerator Laboratory X-ray Free Electron Laser (PAL-XFEL) (Ko *et al.*, 2017, Kang *et al.*, 2017). The photon energy and flux were 9700 eV and 2–5 × 10^11^ photons/pulse, respectively. The X-ray beam size was 8 μm (vertical) × 5 μm (horizontal) (full-width half-maximum) at the sample position, which was focused using a Kirkpatrick-Baez mirror (Kim *et al.*, 2018). The sample holder containing crystals embedded in viscous medium was mounted in the FT-SFX chamber. The raster scanning stage was motioned at 60 μm intervals without synchronization for the arrival FEL pulses. Diffraction data were collected at 23–24°C using an MX225-HS detector (Rayonix, LLC, Evanston, IL, USA: 4 × 4 binding mode) with a 30 Hz readout.

### 2.4. Structure determination

The diffraction patterns were filtered using the *Cheetah* program (Barty *et al.*, 2014) and indexed and scaled using the *CrystFEL* program (White *et al.*, 2016). The phasing problem was solved by molecular replacement with the *MOLREP* program (Vagin & Teplyakov, 2010) with the crystal structures of glucose isomerase (PBD code 5ZYE) (Bae *et al.*, 2018) and lysozyme (PDB code 6IG6) (Park *et al.*, 2019) as the search modes. The model structures were rebuilt and refined using *COOT* (Emsley & Cowtan, 2004) and *PHENIX* (Afonine *et al.*, 2012), respectively. The geometry of the final models structures were analyzed using *MolProbity* (Williams *et al.*, 2018). All figures were generated by *PyMOL* (https://pymol.org/). Data collection and refinement statistics are summarized in Table 1. Coordinates and structure factors have been deposited under accession codes 6LL2 (glucose isomerase embedded in gelatin) and 6LL3 (lysozyme embedded in agarose) in the Protein Data Bank. Diffraction images were deposited under 122 and 123 in CXIDB.

### 2.5. Measurement of background scattering

Background scattering of the gelatin, agarose, polyimide films, and air were measured at the 11C beamline at Pohang Light Source II (Pohang, Republic of Korea) (Park *et al.*, 2017). The X-ray energy and photon flex were 12.659 keV and 1 × 10^11^ photons/s. Gelatin and agarose enclosed in double-side polyimide film (25 μm) had a thickness of less than 300 μm. These material were exposed to the X-ray for 200 ms. Data were collected at 25°C using a Pilatus 6M detector (Dectris, Baden, Switzerland).

## 3. Results and discussion

To develop a sample holder for crystals of various sizes and shapes for FT-SFX experiments, we considered the following criteria for supporting the crystals: (i) The crystal samples should be distributed evenly in the sample holder. (ii) Crystal samples should not sink in the sample holder because of gravity. (iii) The inside of the sample holder should be a hydrated environment to prevent dehydration of the crystalline sample. (iv) The X-ray background scattering from the sample holder material should be negligible for data processing. (v) Crystal samples of random sizes or shapes should be supported in the sample holder. We recently performed SMX experiments using a stable sample delivery method in which viscous medium containing crystal samples were passed through a capillary aligned with the X-rays (Nam, 2020a). In this experiment, the viscous medium prevented the crystal from sinking at a low flow rate by gravity. Based on these results, we considered that that the viscous medium could fix the crystals in the desired space. However, if a crystal sample embedded in the viscous medium is exposed to the atmosphere, the crystals and viscous medium could be dehydrated over time. Dehydration of the crystal sample can alter the crystal space group (Lobley *et al.*, 2016) and result in salt crystal generation from the salt solution in the crystallization solution (Lee *et al.*, 2019). Dehydration of the delivery medium can lead to ring pattern scattering (Park *et al.*, 2019), lowering the quality of the diffraction data and causing physical stress to the crystals. To prevent dehydration from the crystal or delivery medium, the viscous medium containing the crystals was enclosed by a sample holder consisting of two 25 μm polyimide films (Figure 1 and Supplementary Figure S1). Crystal samples supported on silicon, silicon nitride, or nylon-mesh in previously reported sample holders were replaced by viscous medium in this study.

The viscous medium in which crystal samples are embedded produces background scattering when exposed to X-rays, which can affect the signal-to-noise ratio during data processing (Conrad *et al.*, 2015, Nam, 2019). Hydrogel-based viscous delivery medium shows low background scattering (Nam, 2019). In this study, gelatin and agarose were used to support the crystals in the sample holder for FT-SFX, which showed a very low level of background scattering (Nam, 2020a).

We established an approach to support crystal samples in sample holders using gelatin and agarose (Figure 1 and Supplementary Figure S1). After melting the gelatin or agarose materials at >100°C, the gelatin or agarose solution was transferred to a Hamilton syringe and solidified at room temperature. However, when the gelatin or agarose gel was mixed directly with the crystal sample, which caused a physical impact on the crystal sample, the crystal was damaged and diffraction intensity was altered. To overcome this problem, we produced gelatin or agarose fragments by moving the plunger back and forth in the dual syringe setup and passing the sample through the narrow inner diameter of the coupler, as described in the sample preparation of a polyacrylamide injection matrix (Park *et al.*, 2019). Next, the crystal suspension was mixed with the viscous gel fragments in a dual syringe setup. The viscous medium containing crystals was passed through a syringe needle and dispersed and loaded in four regions on a polyimide chip (Figure 1a). Next, the pipette tip was used to disperse the viscous medium containing crystals to spread them evenly over the entire polyimide area (Figure 1b). To prevent dehydration, the sample was immediately covered with another polyimide chip, and the upper and lower polyimide films were enclosed with double-sided adhesive polyimide tape (Figure 1c).

To determine the optimal conditions for supporting the crystals without the effects of gravity when the sample holders were vertically placed, 1–20% (w/v) gelatin or agarose were screened. Gelatin at 10-15% (w/v) and agarose at 1-4% (w/v) were spread evenly on the polyimide film with pipette tips and remained stable between the polyimide films for more than 6 h. However, <8% (w/v) gelatin were displaced by gravity after approximately 10 min in the sample holder. Additionally, >20% (w/v) gelatin and >10% (w/v) agarose gel fragments were highly viscous and rigid, respectively, which physically affected the crystal sample during spreading. Thus, 10% (w/v) gelatin and 4% (w/v) agarose were used for SFX data collection.

To evaluate the viscous medium-based crystal support, we performed FT-SFX experiments at the PAL XFEL. For glucose isomerase embedded in gelatin, 55,307 images were collected for 31 min. The number of hit images containing the diffraction pattern and indexed images were 33,438 (hit rate: 60.45%) and 22,417 (indexing rate: 67.04%), respectively. The data were processed up to 1.75 Å, and overall completeness, signal-to-noise ratio (SNR), R_split,_ and Pearson correlation coefficient (CC) were 100, 5.20, 17.64, and 0.9348, respectively. The R_work_ and R_free_ of the glucose isomerase structure were 17.17 and 19.65, respectively. For lysozyme embedded in agarose, 55,885 images were collected for 31 min. The number of hit images containing the diffraction pattern and indexed images were 24,857 (hit rate: 44.47%) and 14,264 (indexing rate: 57.38%), respectively. The data were processed up to 1.80 Å, and the overall completeness, SNR, R_split,_ and CC were 100, 6.66, 12.43, and 0.9747, respectively. The R_work_ and R_free_ of the final model of lysozyme were 20.27 and 24.62, respectively. Glucose isomerase and lysozyme showed a very clear electron density map for Tyr3-Ala386 and Lys19-Leu147, respectively (Figure 2). No significant radiation damage was observed at the metal binding sites in glucose isomerase and disulfide bonds in lysozyme.

**Figure 2.**
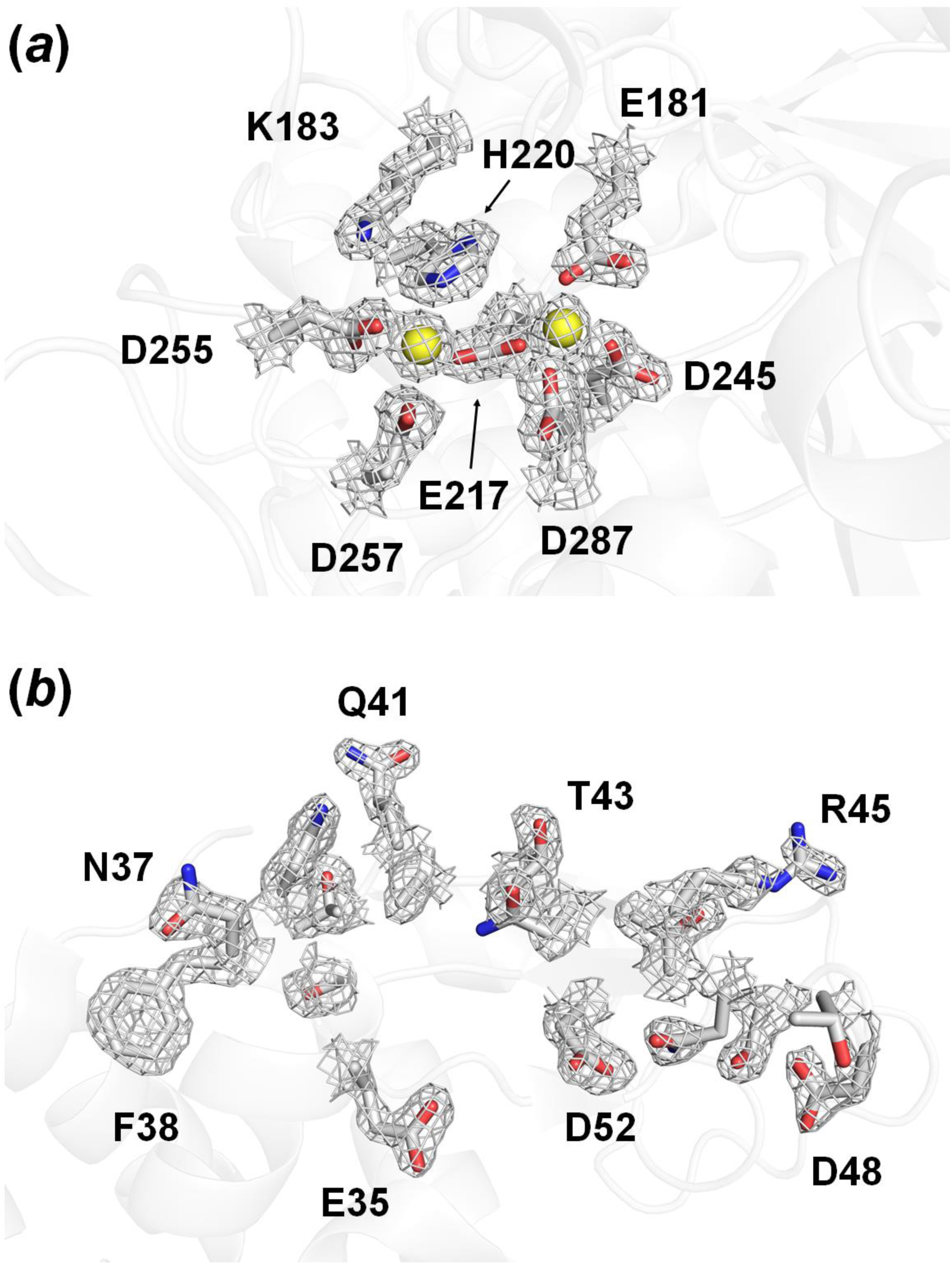
2Fo-Fc electron density map (grey mesh, 1.5 σ) of (a) metal binding site of glucose isomerase embedded in gelatin and (b) active site of lysozyme embedded in agarose.

We monitored the sample holder using a microscope after FT-SFX data collection. As expected, holes were formed in the polyimide film at points where the XFEL beam was transmitted (Figure 3); a similar pattern was observed in previous FT-SFX experiments using mylar film and polyimide film (Doak *et al.*, 2018, Lee *et al.*, 2019). The intervals of the hole positions were 60 μm, which was the interval set for raster scanning at the motion stage. In a previously described nylon-mesh-based sample holder, bubbles considered as radiation damage were observed for each nylon mesh pore created by XFELs (Lee *et al.*, 2019), whereas these bubbles were not observed in the viscous medium inside the polyimide film used in our experiment. We consider that the site at which the XFEL penetrated may have been refilled with viscous medium.

**Figure 3.**
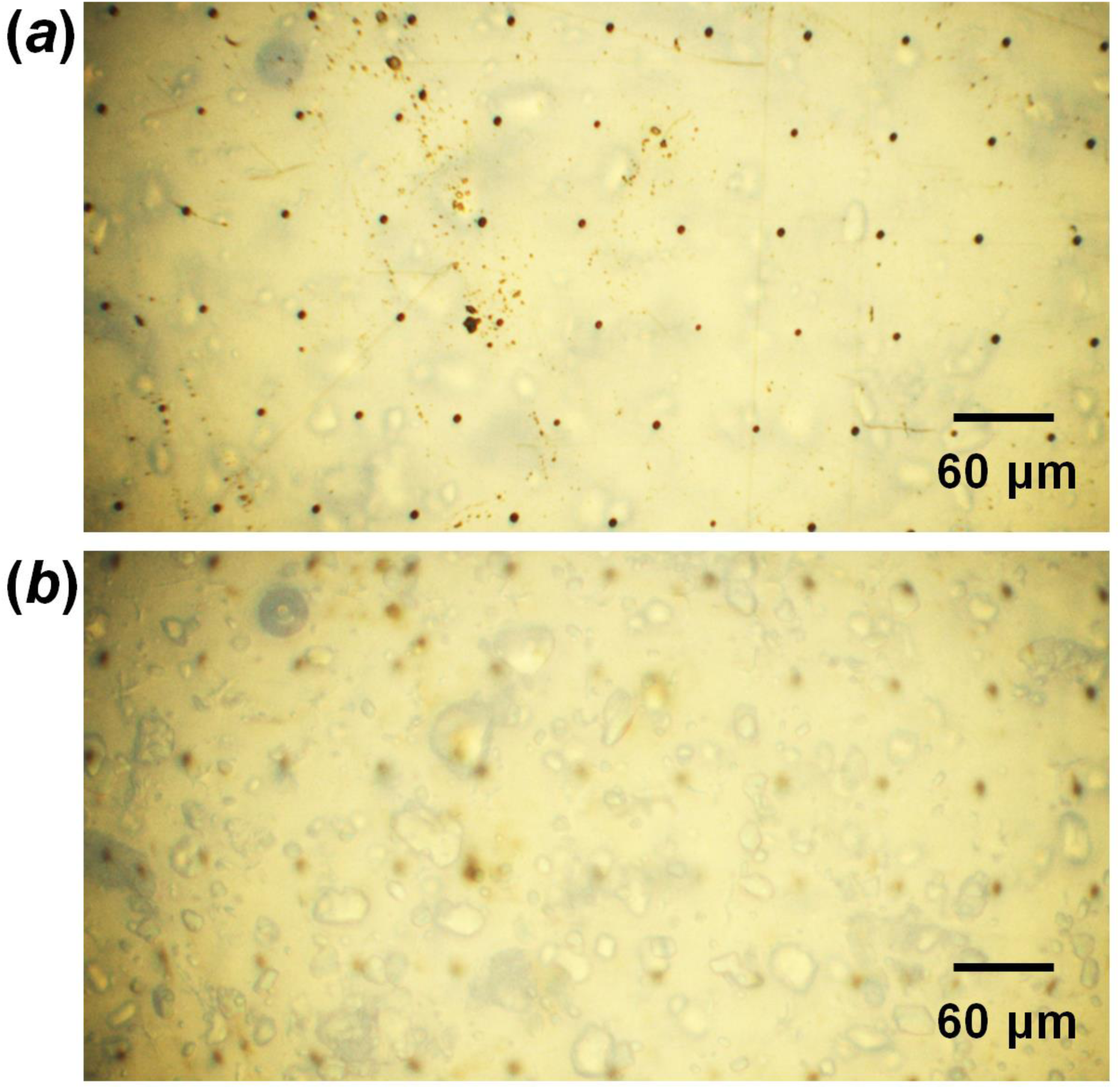
Observation of XFEL penetrated viscous gelatin medium based sample holder. (a) Close-up view of holes generated on the polyimide film through which the XFEL beam was passed. (b) Close-up view of viscous gelatin medium inside of polyimide film through which the XFEL beam was passed.

Next, background scattering of the gelatin and agarose enclosed in the polyimide film, polyimide, and air were measured using a synchrotron X-ray (Figure 4). The thickness of the gelatin and agarose was less than 300 μm when the thickness of polyimide was 25 μm and 2 sheets were used. The total thickness of polyimide is 50 μm. The gelatin and agarose with polyimide film showed intensities of 64.4 and 95.4 analog to digital conversion units (ADU) around the beam stopper, which was considered as background scattering from the delivery medium (Figure 4a and 4b). Diffuse scattering of intensity of 15-29 ADU was observed at approximately 3.2 Å, which was considered to be from solvent. X-Ray background scattering of polyimide was observed at 15.30 Å with 11.6 ADUs (Figure 4c). Air scattering showed values of approximately 2–8 ADU between 5–50 Å (Figure 4d). Therefore, gelatin and agarose with polyimide produced negligible background scattering (Figure 4e).

**Figure 4.**
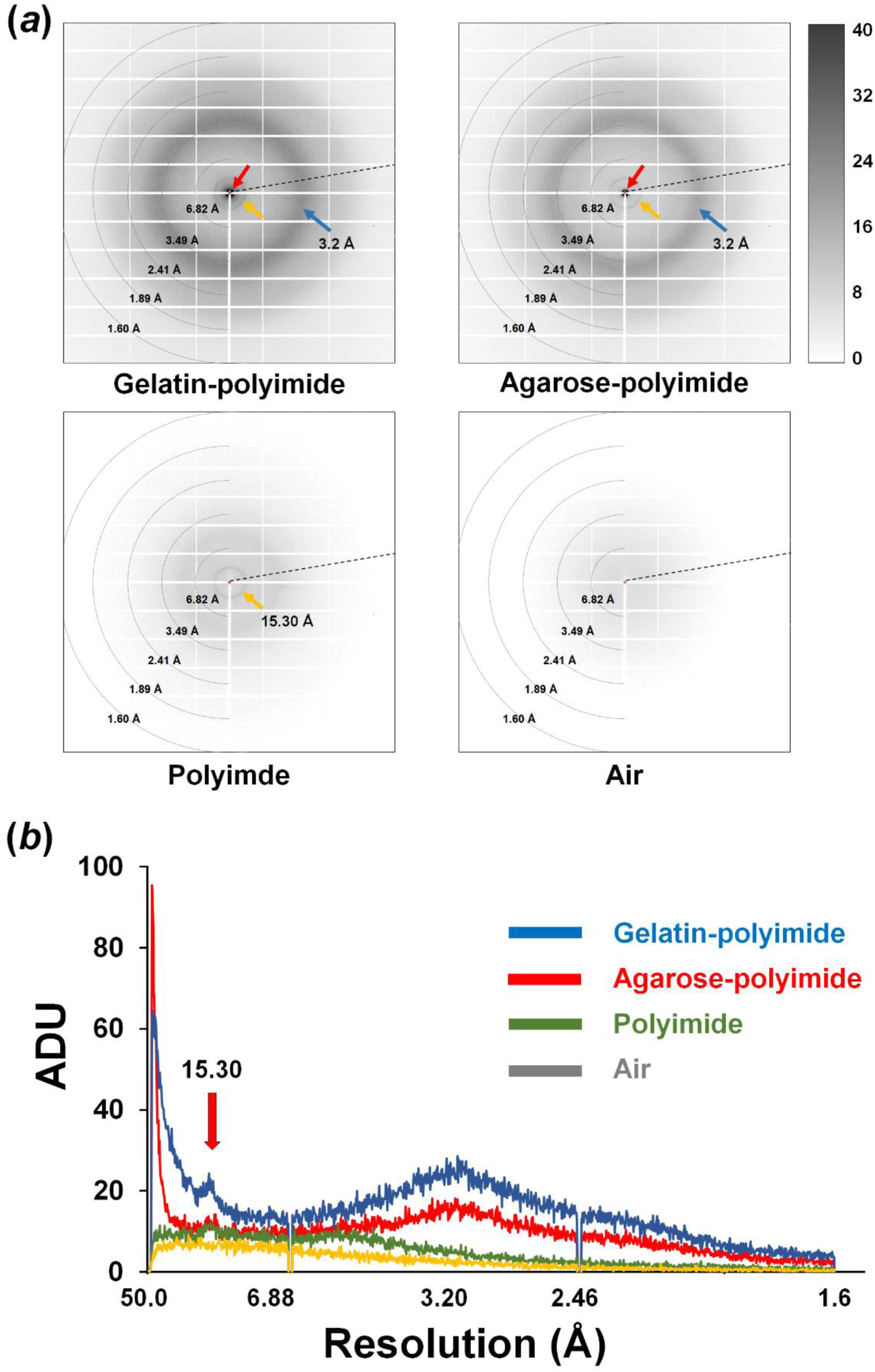
Measurement of background scattering of (a) 300 μm gelatin enclosed with 50 μm polyimide, (b) 300 μm agarose enclosed with 50 μm polyimide, (c) 50 μm polyimide and (d) air. (e) The radial average profile of the scattering intensities of (a)–(d). A scattering ring of polyimide is observed around 15.30 Å.

In summary, we developed a viscous medium-based crystal support for FT-SFX and successfully determined the room temperature structures of glucose isomerase and lysozyme at high resolution. This method can be applied for crystalline samples of various shapes and sizes, and the background scattering generated from the sample holder does not affect data processing. In addition, as the materials constituting the sample holder are composed of transmitted X-rays, the sample holder does not need to be precisely aligned with the X-rays before data collection. This not only simplifies the process but also provides the opportunity to collect more data in limited beamtime. In this experiment, gelatin and agarose were used as the viscous medium, but other viscous delivery media can be applied for crystals supporting. Particularly, selecting a hydrogel-based material with low background scattering would be advantageous for ensuring the quality of data. However, a sample holder based on viscous medium requires mixing of the crystal sample and delivery material, which may not be suitable if large physical damage occurs during mixing. We used a 300 μm thick PVC frame to easy handle the polyimide film in this experiment, and the thickness of viscous medium containing crystals was <300 μm. Although low-level background scattering occurred in the gelatin and agarose medium, a thin viscous medium can avoid X-ray absorption or multiple crystal hits. Therefore, it is important to control the thickness of the viscous medium to the lowest possible level, but consideration should be given to physical damage to the crystal during the thinning process. Our method can be applied for various types of samples in SX studies using XFEL or synchrotron X-ray.

## Acknowledgements

We thank the beamline staff at NCI bemline at Pohang Accelerator Laboratory X-ray Free Electron Laser (PAL-XFEL) for their assistance with data collection. The authors thank Global Science experimental Data hub Center (GSDC) at Korea Institute of Science and Technology Information (KISTI) for computational support.

## Figure legends

Data collection and refinement statistics

| Data collection | Glucose isomerase delivered in gelatin | Lysozyme delivered in agarose |
| --- | --- | --- |
| Energy (eV) | 9700 | 9700 |
| Photons/sec <sup>a</sup> | $\sim 5 \times 10^{11}$ | $\sim 5 \times 10^{11}$ |
| Pulse width | 20 fs | 20 fs |
| Space group | I222 | P4<sub>3</sub>2<sub>1</sub>2 |
| Cell dimensions (Å) |  |  |
| *a* | 92.63 | 79.98 |
| *b* | 98.23 | 79.98 |
| *c* | 101.61 | 38.22 |
| No. collected images | 55307 | 55885 |
| No. of hits | 33438 | 24857 |
| No. of indexed patterns | 22412 | 14264 |
| Resolution (Å) | 70.9-1.75 (1.81-1.75) | 72.4-1.80(1.86 -1.80) |
| Unique reflections | 47004 | 11728 |
| Completeness | 100 (100) | 100 (100) |
| Redundancy | 346.2 (124.5) | 155.1 (63.5) |
| *I*/$\sigma$(*I*) | 5.20 (3.00) | 6.66 (2.06) |
| *R*<sub>split</sub> <sup>b</sup> | 17.64 (23.72) | 12.43 (48.57) |
| CC | 0.9348 (0.7908) | 0.9747 (0.7124) |
| CC* | 0.9830 (0.9397) | 0.9935 (0.9121) |
| Wilson B factor (Å<sup>2</sup>) | 17.91 | 28.95 |
| Refinement statistics |  |  |
| Resolution (Å) | 70.62-1.75 | 55.84-1.80 |
| *R*<sub>factor</sub> / *R*<sub>free</sub> (%) <sup>c</sup> | 17.17 / 19.65 | 20.27 / 24.62 |
| B-factor (Averaged) |  |  |
| Protein | 11.38 | 21.47 |
| Ligands | 5.93 | 30.93 |
| R.m.s. deviations |  |  |
| Bond lengths (Å) | 0.006 | 0.005 |
| Bond angles (°) | 0.831 | 0.725 |
| Ramachandran plot (%) |  |  |
| favoured | 97.12 | 99.21 |
| allowed | 2.62 | 0.79 |
| outliers | 0.26 | 0 |

Highest resolution shell is shown in parentheses.
<sup>a</sup> Sample position
<sup>c</sup> $R_{work} = \sum ||F_{obs}| - |F_{calc}|| / \sum |F_{obs}|$ , where $F_{obs}$ and $F_{calc}$ are the observed and calculated structure-factor amplitudes respectively. $R_{free}$ was calculated as $R_{work}$ using a randomly selected subset (10%) of unique reflections not used for structure refinement.

**Supplementary Figure S1.**
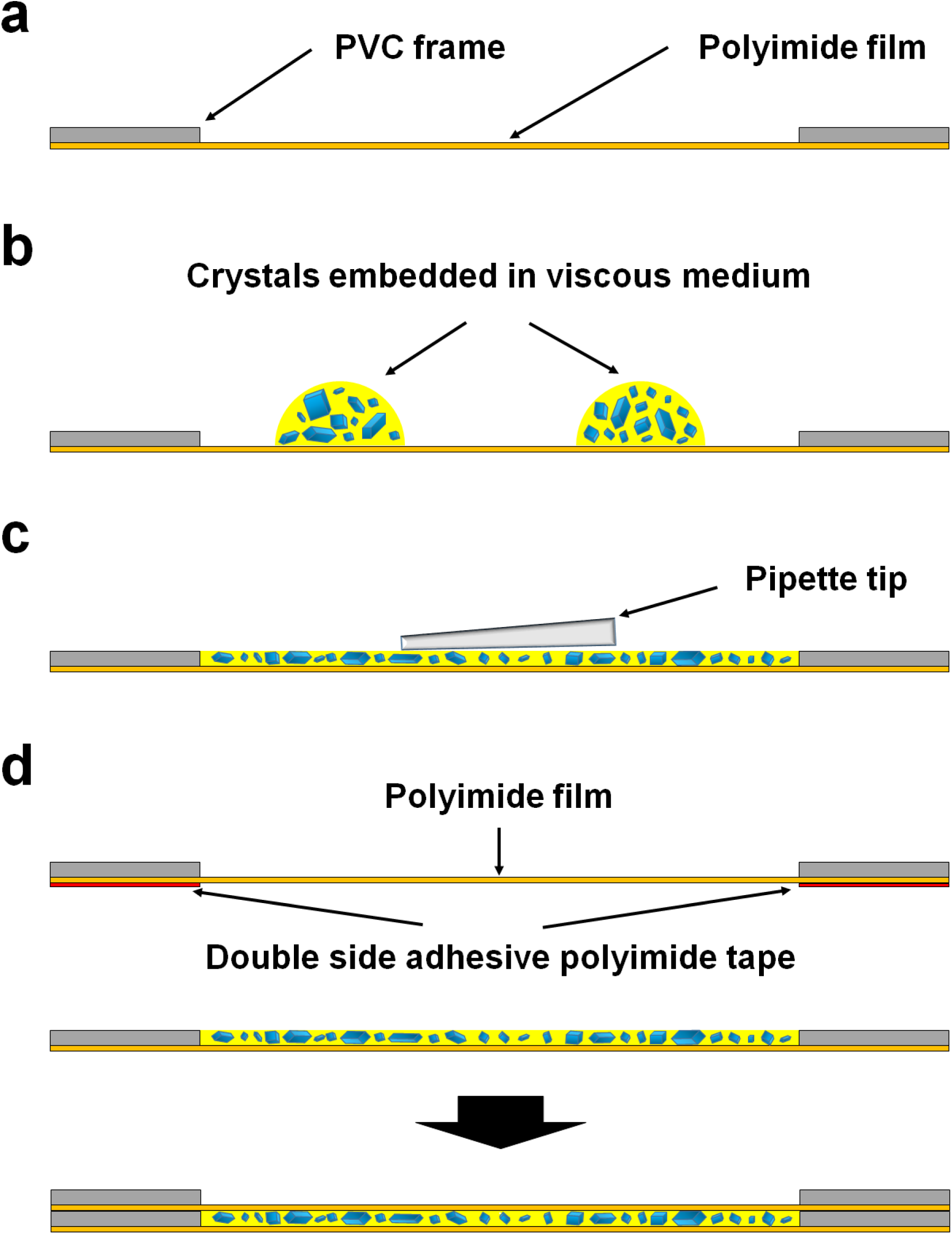
Schematic drawing of sample loading of viscous medium-based sample holder. (a) Sample holder chip consisting of polyimide film and PVC frame. (b) Loading of crystals embedded in viscous medium using a pipette tip. (c) Sample spreading using a pipette tip. (d) Covering of the sample using another sample holder chip.

